# Mice identify subgoal locations through an action-driven mapping process

**DOI:** 10.1101/2021.12.14.472688

**Authors:** Philip Shamash, Tiago Branco

## Abstract

Mammals instinctively explore and form mental maps of their spatial environments. Models of cognitive mapping in neuroscience mostly depict map-learning as a process of random or biased diffusion. In practice, however, animals explore spaces using structured, purposeful, sensory-guided actions. Here we test the hypothesis that executing specific exploratory actions is a key strategy for building a cognitive map. Previous work has shown that in arenas with obstacles and a shelter, mice spontaneously learn efficient multi-step escape routes by memorizing allocentric subgoal locations. We thus used threat-evoked escape to probe the relationship between ethological exploratory behavior and allocentric spatial memory. Using closed-loop neural manipulations to interrupt running movements during exploration, we found that blocking runs targeting an obstacle edge abolished subgoal learning. In contrast, blocking other movements while sparing edge-directed runs had no effect on memorizing subgoals. Finally, spatial analyses suggest that the decision to use a subgoal during escape takes into account the mouse’s starting position relative to the layout of the environment. We conclude that mice use an action-driven learning process to identify subgoals and that these subgoals are then integrated into a map-based planning process. We suggest a conceptual framework for spatial learning that is compatible with the successor representation from reinforcement learning and sensorimotor enactivism from cognitive science.

## Introduction

A fundamental ability of mobile animals is to learn the location of resources and how to get there. This can in principle be done using a variety of strategies. At one end, the behaviorist framework focuses on the importance of repeating actions. Mazes can be solved, after sufficient practice, by learning the correct movements directly in a ‘stimulus-response sequence’ (Hull 1934; Restle 1957a). At the opposite end, the cognitive map theory proposes that animals possess mental maps of their environments that they can query to navigate to goals (Tolman 1948). In this framework, a spatial map is learned through an innate capacity to map observations and is used to derive novel actions (O’Keefe and Nadel 1978). These two learning strategies are thought of as independent processes within the brain: a striatal system for repeating successful movements and targeting visible landmarks; and a hippocampal system for constructing an internal map of the environment (Doeller et al. 2008; Packard et al. 1989).

Cognitive maps are particularly powerful because they decouple actions from spatial learning, allowing the computation of routes in an allocentric (spatial-location-centered) reference frame. Models of this class therefore do not generally consider the motivation underlying the learner’s exploratory actions; instead, they use ‘random agents’ that repeatedly select movements from a distribution of directions and distances in order to map out as many locations as possible (e.g. Burgess et al. 1994; Stachenfeld et al. 2017; Viswanathan et al. 1999; but see McNamee et al. 2021). Similarly, the paradigmatic experimental studies in this vein focus on the cues rather than the actions that animals use to pinpoint locations, and they employ a session structure that ends immediately after the animal locates the reward (Cheng et al. 2013; Morris 1981; Restle 1957a; Tolman and Honzik 1930). These methodologies contrast starkly with the way animals explore natural environments. Mice, for example, move about in a highly structured manner, punctuating investigatory bouts along boundaries with rapid lunges to a familiar, enclosed space or a visually salient object (Crowcroft 1966). It thus seems plausible that the sensorimotor tendencies of each species could play an important role in identifying important locations or compartments within the map, rather than serving a fully independent function (Alyan 2004; Ballard et al. 1997; Clark 1999; Mataric 1992).

The homing behavior of rodents offers a powerful window into the relationship between spontaneous exploration patterns and spatial cognition (Evans et al. 2019). Within minutes of entering a new environment, rodents rapidly identify and memorize sheltering locations (Vale et al. 2017); spontaneously shuttle back and forth between the outside and the ‘home’ (Crowcroft 1966; Maaswinkel and Whishaw 1999; Shamash et al. 2021); and respond to threatening stimuli by running directly to shelter (Yilmaz and Meister 2013). Moreover, homing behavior is sophisticated enough to involve map-based computations of multi-step escape routes. Shamash et al. 2021 recently showed that mice escape past obstacles by memorizing allocentric subgoal locations at the edges of the obstacle. Intriguingly, the learning of subgoal locations was highly correlated with the execution of a particular sensorimotor action during the exploration period - spontaneous running movements targeting the obstacle edge. This raises the hypothesis that the execution of specific exploratory actions is important for learning elements of a cognitive map.

Here we directly test this hypothesis by investigating whether spontaneous edge-directed runs are causally relevant for subgoal learning. We use closed-loop neural manipulations to precisely interrupt these runs during exploration, and then examine the effect on the use of subgoals during escape behavior. We demonstrate that subgoal learning is action-driven in nature, and then go on to show that it also relies on a mapping capacity. We suggest that spatial learning through natural exploration relies on a learning mechanism that combines both action- and map-based strategies.

## Results

### Closed-loop optogenetic activation of premotor cortex to block spontaneous edge-vector runs

When mice are placed in an arena with a shelter and an obstacle, they spontaneously execute runs targeting the obstacle edge (Shamash et al. 2021). Our main aim in this work was to test the causal necessity of these runs in learning that the obstacle edge is a subgoal, i.e. a location that should be targeted when attempting to run past the obstacle to get to the shelter. We therefore designed a manipulation that could prevent mice from executing spontaneous runs to an obstacle edge. To prevent confounding effects, our manipulation should also avoid modifying the external environment, should not decrease the opportunities for the animal to observe its environment, and should not generate place aversion. We found that closed-loop stimulation of premotor cortex (M2) fit all these criteria. We expressed channelrhodopsin in excitatory neurons in the right, anterior M2, and performed optogenetic stimulation via an implanted optic fiber (Fig. 1b, Supp. Fig. 1a). In line with previous reports (Gradinaru et al. 2007; Magno et al. 2019), stimulating M2 with a 2-sec, 20-Hz pulse wave caused a low-latency (<200 ms) deceleration, halting, and leftward turning motion (Supp. Fig. 1b; Video 1). This stimulation protocol did not generate place aversion when tested in a two-chamber place-preference assay (Supp. Fig. 1d). We thus leveraged this approach to specifically interrupt edge-vector runs during spontaneous exploration. Using online video tracking, we set up a virtual “trip wire” in between the threat area and the left obstacle edge; whenever mice crossed this line while moving in the direction of the edge, a 2-sec pulse of light was automatically delivered (Fig. 1c; Video 1). Up to three subsequent pulses were triggered manually if the mouse continued moving toward the edge. All other movements, including runs to the left edge along the obstacle or from the shelter, were not interrupted by laser stimulation.

**Figure 1:**
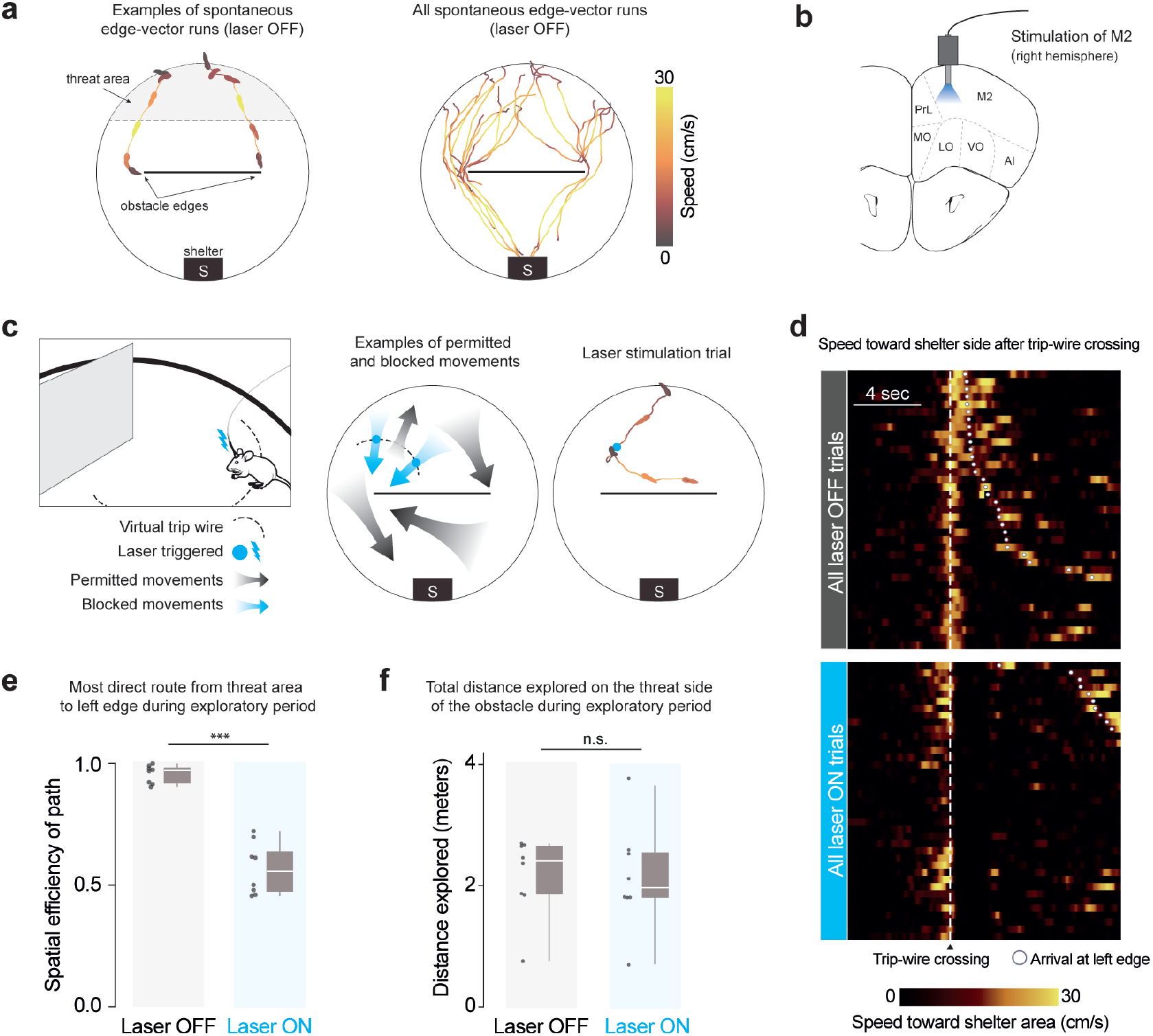
Closed-loop optogenetic activation of M2 interrupts spontaneous edge-vector runs **(a)** Spontaneous edge-vector runs are continuous turn-and-run movements, starting in the threat area and stopping at or moving past the obstacle edge, during the initial 20-minute exploration period. n=8 mice. **(b)** Optic fibers were implanted in right premotor cortex, .25 mm above the channelrhodopsin injection site. M2: supplementary motor cortex (premotor cortex), PrL: prelimbic cortex, MO/LO/VO: medial/lateral/ventral orbital cortex, AI: agranular insular cortex. **(c)** On crossing an invisible trip wire (dotted line) during exploration, mice automatically received a 2-sec, 20-Hz, 30-mW pulse of 473-nm light. This caused a stopping and leftward-turning motion. Up to three subsequent 2-sec pulses were triggered manually if the mouse continued moving forward. In the example trial, the mouse was stimulated with two 2-sec pulses and then ran to the right side of the platform. Mouse drawing: scidraw.io. **(d)** All trip-wire crossings, with and without laser stimulation, ordered by time of arrival to the left obstacle edge. Note that mice must be moving toward the shelter area (i.e., southward) in order to trigger the trip wire. **(e)** Spatial efficiency is the ratio of the straight-line path to the length of the path actually taken. White horizontal lines indicate median, gray boxes indicate the first and third quartiles, and gray vertical lines indicate the range. Each dot represents one mouse/session. p=5*×* 10^−5^, one-tailed permutation test. **(f)** Distance explored on the threat half: p=0.5, one-tailed permutation test. n=8 mice in each group.

We divided up injected and implanted animals into a laser-on (trip wire active) and a control, laser-off group (trip wire inactive). Both groups of mice were allowed to explore a circular platform with a shelter and an obstacle for 20 minutes (n=8 mice/sessions; pictured in Supp. Fig. 2). During this time, all mice located the shelter and visited the entire platform, including the obstacle (Supp. Fig. 3a,c). In agreement with previous results (Shamash et al. 2021), all mice in the laser-off group executed continuous running movements from the threat area (Fig. 1a) toward the shelter area (‘homing runs’; # per session: 6 [5, 8.25] (median [IQR]); Methods; Supp. Fig. 3b,d). These included at least one homing run that directly targeted an obstacle edge (‘edge-vector runs’; # per session: 1.5 [1, 2.25] (median [IQR]); Fig. 1a, Supp. Fig. 3e; Video 1). Mice in the laser-on group triggered 3.5 [2.75,6] (median [IQR]) laser stimulation trials, lasting 20 [16, 26] seconds in total and interrupting all potential edge-vector runs (Fig. 1d, Supp. Fig. 3b,e). While mice in the laser-off group executed nearly direct paths between the threat area and the left obstacle edge, the paths taken by mice in the stimulation group were twice as long, reflecting the inaccessibility of edge-vector runs (Fig. 1e). Exploratory behavior in general, however, was not reduced. Mice in the stimulation condition explored the obstacle, the edge, the threat area and the entire arena as much as the control group (Fig. 1e, Supp. Fig. 3a,c).

**Figure 2:**
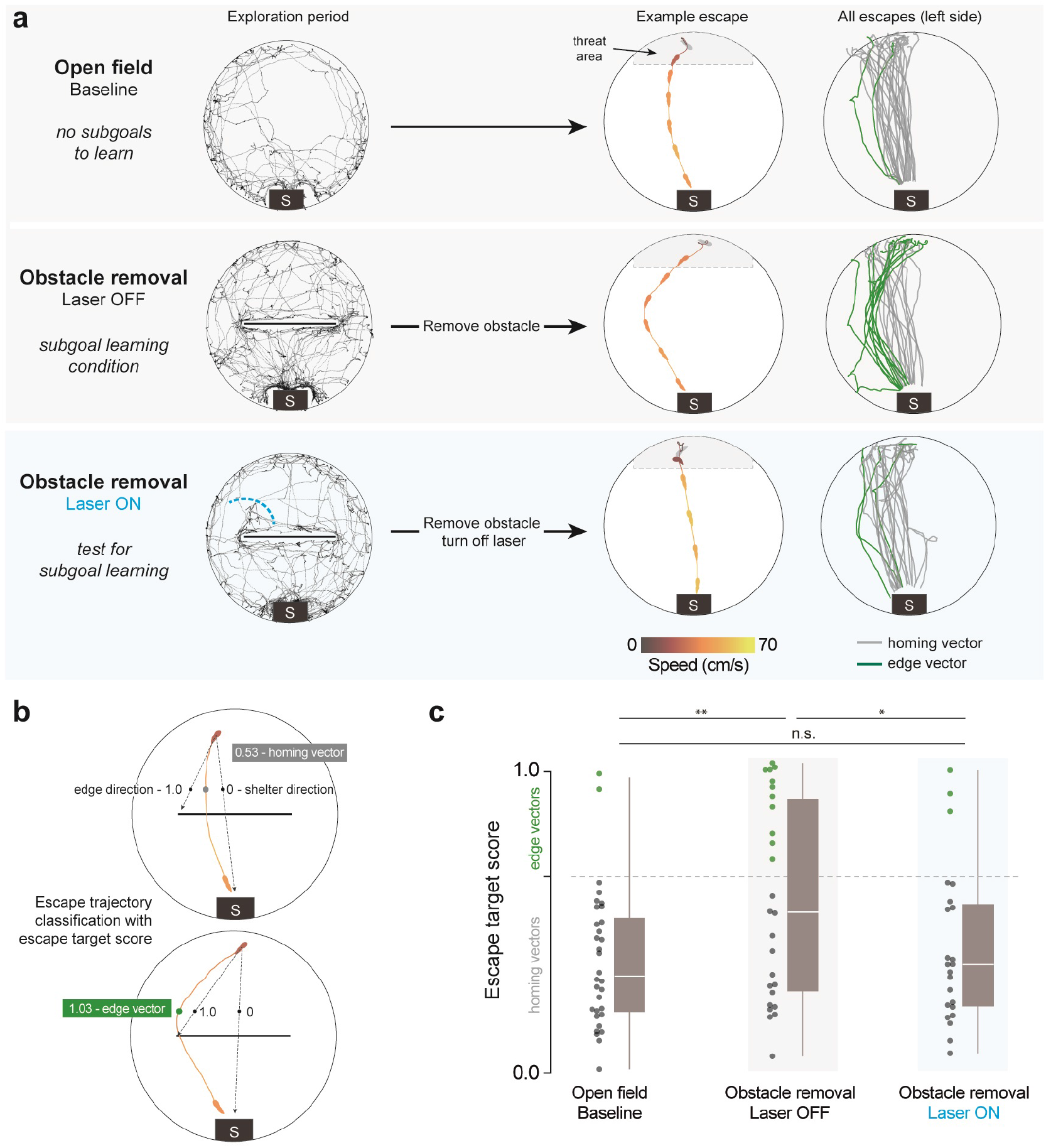
Interrupting spontaneous edge-vector runs abolishes subgoal learning **(a)** Black traces show exploration during an example session (open field: 10 mins, obstacle removal: 20 mins). After this, escapes are triggered automatically in the threat zone. Lines and silhouette traces show escape routes from threat onset to shelter arrival. Since we are examining the effect of blocking runs toward the left obstacle edge, we limited analysis to escapes on the left side of the platform (Supp. Fig. 4b). Open field: 29 escapes; Obstacle removal (laser off): 26 escapes; Obstacle removal (laser on): 23 escapes. All: n=8 mice. **(b)** The initial escape target is the vector from escape initiation to 10 cm in front of the obstacle (black dots), normalized between 0 (shelter direction) and 1 (obstacle edge direction). Escape initiation is where the mouse’s speed relative to the shelter exceeds 20 cm/sec. **(c)** As in Shamash et al. 2021, escape target scores over 0.65 are classified as edge vectors; scores under 0.65 are classified as homing vectors. Obstacle removal (laser off) vs. open field: p=.003; Obstacle removal (laser on) vs. open field: p=.2; Obstacle removal (laser off) vs. obstacle removal (laser on): p=.03, one-tailed permutation tests on proportion of edge-vector escapes.

**Figure 3:**
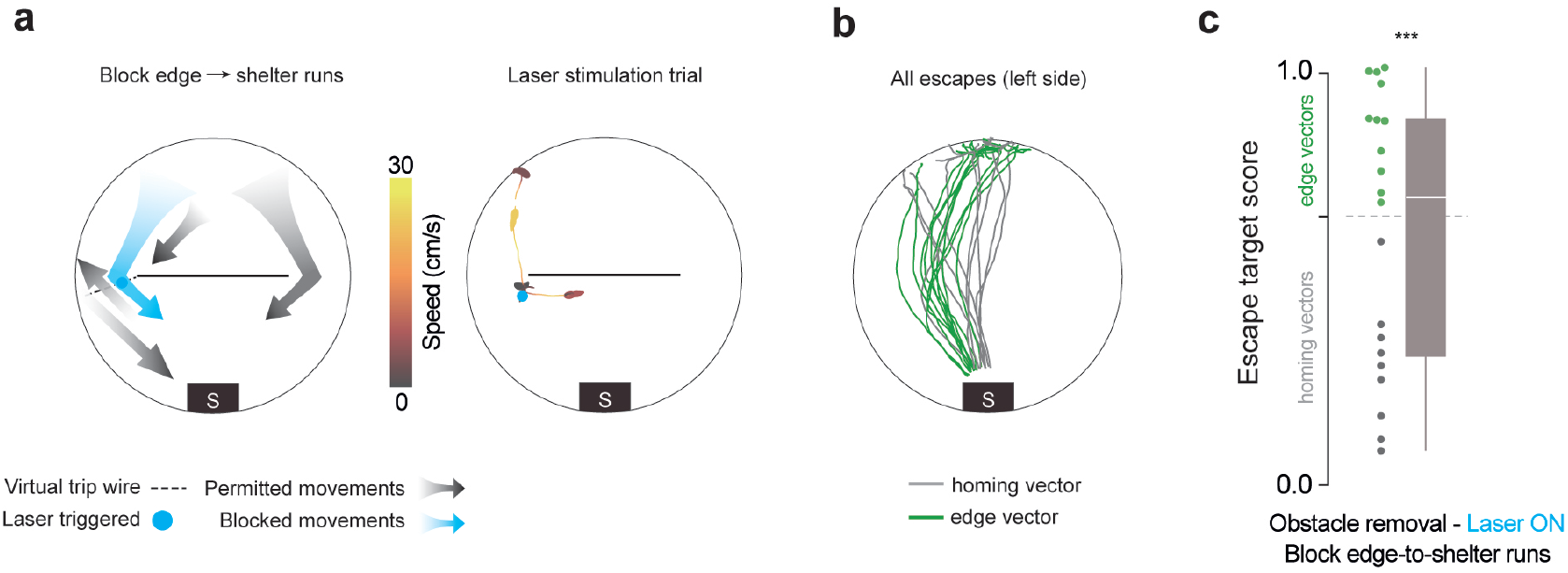
Blocking edge-to-shelter runs does not diminish subgoal learning **(a)** Blocking left-edge-to-shelter runs. Instead of one-four 2-sec laser pulses, here we delivered one-two 5-sec pulses. This longer duration served to keep the mouse from getting to the shelter for longer. In the example trial, the mouse was stimulated for ten seconds, and then ran toward the center of the platform. **(b)** Escapes after obstacle removal. n=8 mice, 23 escapes (left side). **(c)** Obstacle removal (block edge-to-shelter) vs. open field: p=1 *×*10^−4^ (***); vs. obstacle removal (block edge vectors): p=.03; vs. obstacle removal (laser off): p=.8; one-tailed permutation tests on proportion of edge-vector escapes.

### Interrupting spontaneous edge-vector runs abolishes subgoal learning

We next measured the impact of blocking edge-vector runs on subgoal learning. After the 20 min exploration period, we elicited escape behavior using a loud, unexpected crashing sound. Mice triggered an auditory threat stimulus automatically by entering the threat zone and staying there for 1.5 seconds. Escape routes were quantified using a target score and classified as targeting the obstacle edge (‘edge vector’) or the shelter (‘homing vector’) (Fig. 2b; see Methods).

First, we acquired a negative-control distribution by letting a group of mice explore and escape in an open-field environment with no obstacle (n=8 mice; same viral injection and implantation procedure as above). As expected from previous work (Vale et al. 2017), mice generally responded to threats by turning and running directly along the homing vector (Fig. 2a, Supp. Fig. 4a; Video 2). Second, we examined escapes in a positive-control condition known to generate subgoal learning. After the laser-off group explored the arena with the obstacle and shelter for 20 minutes, we removed the obstacle and triggered escapes (2-30 minutes later, IQR: 8-17 minutes). We found that 42% of escapes were directed toward the obstacle edge location, despite the obstacle being gone (‘edge vectors’; 26 total escapes on the left side; more edge vectors than in the open field: p=0.003, permutation test; Fig. 2a,c; right-side escapes in Supp. Fig. 4b; Video 3). This result is consistent with Shamash et al. 2021, which found that these edge-vector escapes reflect the memorization of a subgoal location.

**Figure 4:**
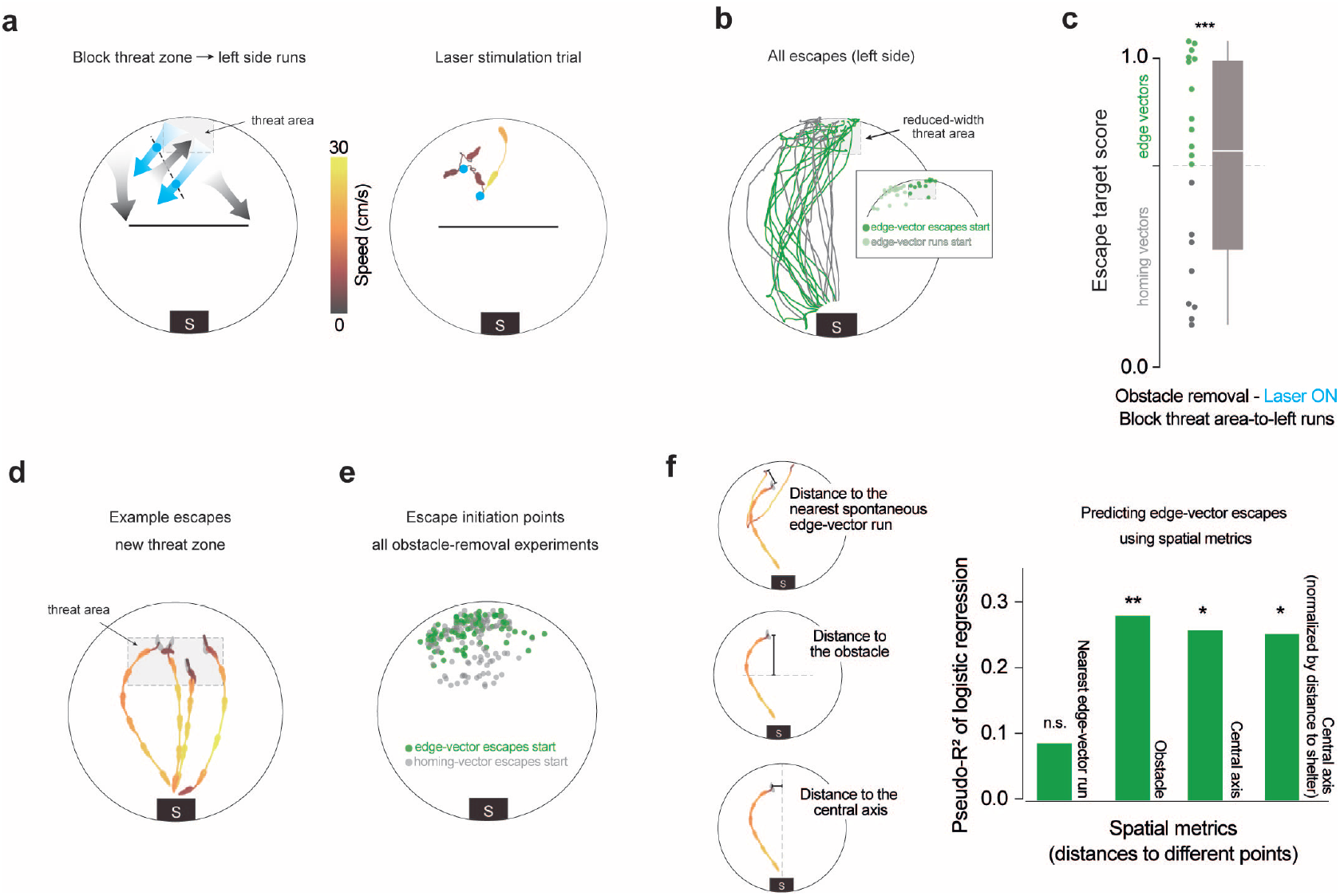
Subgoal-escape start points are determined by spatial rules, not edge-vector-run start points **(a)** Blocking threat-zone-to-left-side runs. The stimulation protocol is the same as in figure 1, except with a new trip-wire location. The dotted red line outlines the threat zone used in this experiment. In the example trial, there were two consecutive trip-wire crossings (2-sec stimulations), after which the mouse moved back toward the threat zone. **(b)** Escapes after obstacle removal. The reduced-width threat zone ensured that mice would need to cross the deactivated trip-wire in order to execute edge-vector escapes. n=8 mice, 19 escapes (left side). Inset: All start locations for spontaneous edge-vector runs (light green) and subsequent edge-vector escapes (dark green). **(c)** Obstacle removal (block threat-zone-to-left-side) vs. open field: p=6*×* 10^−4^ (***); vs. obstacle removal (block edge vectors): p=.01; vs. obstacle removal (laser off): p=.8, one-tailed permutation tests on proportion of edge-vector escapes. **(d)** Four example escapes triggered after obstacle removal, using the new threat zone. **(e)** Data from all obstacle-removal experiments in this paper are combined here (with the exception of the block-edge-vectors experiment). These include escapes on both the left and right sides. In order to avoid uncertainty over which edge the edge-vector escapes are targeting, right-sided escapes are flipped horizontally in this visualization; thus, all the green dots can be seen as left-edge vectors. Each dot represents one escape. n=40 sessions, 207 escapes. **(f)** McFadden’s pseudo-R^2^ measures the strength of the relationship between each metric and the odds of executing edge-vector escapes. Values from 0.2 to 0.4 represent “excellent fit” (McFadden 1977). Distances are measured from the escape initiation point of each escape. For the distance to the nearest spontaneous edge-vector run start point, only runs toward the same side as the escape are considered. Distance to the nearest start point of a spontaneous edge-vector run: pseudo-R^2^=0.086; p=0.5. Distance to the obstacle: pseudo-R^2^=0.28; p=0.007. Distance to the central axis: pseudo-R^2^=0.26; p=0.01. Normalized distance to the central axis: pseudo-R^2^=0.25; p=0.02. P-values come from a permutation test using 10,000 random shuffles of the edge-vector/homing-vector labels, with the pseudo-R^2^ as the test statistic.

Third, we tested the laser-on group, which explored with an obstacle and shelter but had their exploratory edge-vector runs interrupted. After removing the obstacle, threat-evoked escape routes resembled the paths taken in the open-field condition rather than the subgoal-learning group (13% edge vectors; 23 escapes (left side); fewer edge vectors than in the laser-off condition: p=0.03, and not significantly more edge vectors than in the open field: p=0.2, permutation tests; Fig. 2a,c). Thus, interrupting spontaneous edge-vector runs abolished subgoal learning.

An alternative explanation could be that these mice did learn subgoals, but the stimulation during edge-vector runs taught them to avoid expressing edge-vector escapes. To address this possibility, we repeated the stimulation experiment (n=8 mice), this time allowing mice to perform two spontaneous trip-wire crossings without interruption. We then subjected them to the same edge-vector-blocking protocol as above (3 [1.75, 4.25] laser trials per session (median [IQR]) lasting 16 [5.5, 26.5] secs in total; Supp. Fig. 5a, 6; Video 4). Removing the obstacle and triggering escapes now revealed robust subgoal behavior (65% edge vectors; n=23 escapes (left side); more edge vectors than in the open field: p=3*×* 10^−4^, and not significantly fewer edge vectors than the laser-off condition: p=.9, permutation tests). This shows that our manipulation does not reduce the use of subgoals once they are learned and therefore suggests that edge-vector runs are causally required for learning subgoals.

### Blocking edge-to-shelter runs does not diminish subgoal learning

Spontaneous edge-vector runs are often followed by an edge-to-shelter run. After completing an edge-vector run, mice in the laser-off condition reach the shelter within 2.5 [1.7,10] secs (median [IQR]), generally taking direct paths (spatial efficiency: .87 [.47, .95]; 1.0 corresponds to the direct path; Supp. Fig. 6c,e). We therefore considered whether edge-vector runs support subgoal learning because they are part of a sequence of actions that quickly brings the mouse from the threat zone to the shelter.

To test whether edge-to-shelter runs are important for learning, we repeated the stimulation experiment (n=8 mice), but with a new trip-wire location. Using 10-sec laser pulses, we stopped movements from the left obstacle edge toward the shelter (restricted to edge-to-shelter movements that occurred after having crossed the original trip wire, i.e. the second phase of a threat-area-to-edge-to-shelter run; 3 [2, 3.25] laser trials per session (median [IQR]) lasting 25 [20, 30] secs in total; Fig. 3a; Supp. Fig. 6; Video 4). Due to this manipulation, edge-vector runs on the left side were followed by long, slow paths to shelter (seconds to shelter: 29 [18, 55]; spatial efficiency: .28 [.13, .37]; slower than the laser-off condition: p=1*×* 10^−3^; less spatially efficient than the laser-off condition: p=2*×* 10^−3^, permutation tests; Supp. Fig. 6c,e). Despite this effect, removing the obstacle and triggering escapes revealed robust subgoal behavior (55% edge vectors; n=23 escapes (left side); Fig. 3b-c; more edge vectors than in the open field: p=1*×* 10^−4^, and not significantly fewer edge vectors than the laser-off condition: p=.8, permutation tests). Thus, for their causal role in subgoal learning, edge-vector runs do not need to be rapidly followed by the extrinsic reward of entering the shelter. This result also supports the argument that optogenetic stimulation at the left edge does not teach the mice to avoid passing by that location during escapes.

### Subgoal-escape start points are determined by spatial rules

The results from the previous experiment suggest that learning subgoals with edge-vector runs is not simply a matter of reinforcing actions that lead to the shelter. This fits with the finding in Shamash et al. 2021 that subgoals in this context are stored as allocentric locations rather than egocentric movements, and it raises the possibility that the learning process combines actions and spatial information. To explore this further, we investigated the rules governing the set of locations from which mice initiate memory-guided subgoal escapes - the “initiation set” of subgoal escapes. We aimed to determine whether the initiation set is 1) spread indiscriminately throughout the environment; 2) restricted to the vicinity of previous edge-vector-run start positions; or 3) related to the spatial layout of the environment, independent of past actions. Options 1 and 2 would be expected if mice were merely learning to repeat edge-vector actions, while option 3 would be expected if subgoals were embedded in map-based planning process. We first repeated the obstacle removal experiment but now elicited escapes from in front of the obstacle location, near to the shelter (n=8 mice with no laser stimulation, 28 escapes; Supp. Fig. 7a). From this starting point, mice did not escape by running toward a subgoal location but instead fled directly to shelter. This result suggests that the initiation set is spatially confined rather than indiscriminate.

Next, we tested whether the initiation set is confined to the area in which spontaneous edge-vector homing runs had previously occurred. We modified our laser stimulation experiment with a new trip wire location, so that edge-vector runs were allowed from a section of the arena next to the threat zone, but were interrupted if they started within the threat zone (n=8 mice; 2 [1.75, 4] laser trials per session (median [IQR]) lasting 4 [6, 9] secs; Fig. 4a,b; Supp. Fig. 6; Video 4). As before, laser stimulation succeeded in blocking edge-vector runs from the threat zone (Supp. Fig. 6f). In this configuration, however, mice were still able to execute edge-vector runs starting from the area to the left of the threat zone (illustrated by the leftmost gray arrow in Fig. 4a; Supp. Fig. 6d). Removing the obstacle and triggering escapes in this cohort revealed robust subgoal behavior (63% edge vectors; n=19 escapes (left side); Fig. 4b-c; more edge vectors than in the open field: p=6 *×*10^−4^, and not significantly fewer edge vectors than the laser-off condition: p=.8, permutation tests). Thus, the initiation set for subgoal escapes extends beyond the locations in which successful edge-vector runs have been initiated (Fig. 4b inset). This result also reaffirms that optogenetic stimulation does not teach mice to avoid paths that are blocked by laser stimulation during exploration.

To more precisely examine the impact of spatial location on subgoal behavior, we repeated the obstacle removal experiment with a larger threat zone, located between the obstacle location and the original threat zone (n=8 mice, 53 escapes; no laser stimulation; Fig. 4d, Supp. Fig. 7b). By combining these escapes with the original threat zone data, we could test the relationship between the location of escape onset and the tendency to use a subgoal, using logistic regression (n=40 total sessions, 207 escapes; Fig. 4e-f). We found that being closer to previous edge-vector runs was not related to the likelihood of executing edge-vector escapes (McFadden’s pseudo-R^2^=0.086; p=0.5, permutation test; Fig. 4f, Supp. Fig. 7c-d). In contrast, a number of spatial metrics were effective predictors of edge-vector escape probability (Fig. 4f, Supp. Fig. 7c-e). These include the distance from the obstacle, the distance from the central axis of the platform (the axis perpendicular to the obstacle), the distance from the shelter, and the angle between the edge-vector and homing-vector paths. Since these metrics are correlated with each other, we analyzed whether their relationship to escape behavior could all be parsimoniously explained by a sense of distance from the shelter. To test this, we normalized the mouse’s distance from the central axis such that it was uncorrelated to the distance from the shelter, using linear regression. This normalized spatial metric retained its capacity to predict edge-vector escapes (pseudo-R^2^=0.25; p=0.017, permutation test; Fig. 4f). This means that, at a given distance from the shelter, mice are more likely to execute edge-vector escapes if they are further from the central axis of the arena (Supp. Fig. 7d). Thus, mice appear to be keeping track of their two-dimensional position within the arena - their distance from the shelter and obstacle as well as their position along the left-right axis - and using this information to select whether to execute a subgoal-based or homing-vector escape. Overall, our results indicate that the initiation set is defined in relation to the spatial layout of the environment rather than proximity to previous successful actions.

## Discussion

When a mouse investigates a new environment, it does not act like a ‘random agent’. Instead, its exploration consists of purposive, extended, sensorimotor actions. In this work, we have demonstrated that one such class of movements - running to an obstacle edge that grants direct access to a goal - plays a causal role in the process of gaining useful spatial information about the environment.

In our previous work we found that, during 20 minutes of exploration with a shelter and an obstacle, mice memorize subgoals at the obstacle edge location (Shamash et al. 2021). This is revealed by removing the obstacle and presenting threats, which causes mice to initiate escapes by running to the location of an edge that is no longer there. To explain an allocentric behavior such as this, typical spatial learning models would rely on two steps: 1) constructing an internal map of space by observing how locations and obstructions in the environment are positioned relative to each other; and 2) using this map to derive a useful subgoal location, computed either at decision time or in advance during rest (Edvardsen et al. 2020; Spiers and Gilbert 2015) This process is well suited for agents that learn by diffusing throughout their environment, be it randomly or with a bias toward unexplored territory (Schulz and Gershman 2019). However, it does not account for the prevalence of goal- and object-oriented actions in natural exploratory patterns (Crowcroft 1966; Schulz et al. 2017). We thus explored a potential role for a third process: 3) executing ‘practice runs’ to candidate subgoal locations during exploration. We tested this idea using closed-loop optogenetic stimulation of M2 to interrupt spontaneous edge-vector homing runs. This manipulation abolished subsequent subgoal behavior. An important point from the outset is that this effect does not suggest that M2 is important for computing subgoals; three other M2 stimulation protocol that spared edge-vector runs failed to have an effect on learning. Instead, it demonstrates that the edge-vector runs themselves are causally necessary for triggering subgoal memorization in this setting.

One interpretation of our results could be that subgoal behavior is a naturalistic form of operant conditioning; practice edge-vector runs are followed by reinforcement, and then get repeated in response to threat. This framework could explain why edge-vector responses persist after obstacle removal: they are habits that have not yet been ‘extinguished’. Moreover, the lack of effect of blocking edge-to-shelter runs fits with an instrumental chaining mechanism (Gollub 1977; Hull 1934), in which arrival at the obstacle edge itself acts as a reinforcer. On the other hand, subgoal learning diverges from instrumental learning in two ways: it operates within an allocentric framework (generally seen as distinct from an instrumental response strategy (Doeller et al. 2008; Geerts et al. 2020; Packard et al. 1989; Restle 1957b), and it only requires 1-2 practice runs (even simple instrumental training takes tens of learning trials (Baron and Meltzer 2001)). More importantly, the set of locations from which mice initiate subgoal escapes are defined by the mouse’s spatial position relative to the obstacle and shelter, and not by their proximity to previous edge-vector runs. The concepts of action and reinforcement are therefore insufficient for explaining subgoal memorization; planning with an internal map of space must also be invoked.

In line with our results, the successor representation (SR) is a model-based/model-free hybrid mechanism in reinforcement learning (RL) that can achieve map-like planning while also taking into account the speed and direction of exploratory actions (Dayan 1993). In SR planning, RL agents build a representation of where they are likely to go in the future, given a starting location; this is updated after each movement. They can then use a low-dimensional representation of this ‘predictive map’ to identify subgoal locations (Stachenfeld et al. 2017). Alternatively, a predictive-map variant called the first-occupancy representation (FR) natively allows for subgoal planning (Moskovitz et al. 2021). These subgoal-identification processes align with our results demonstrating the necessity of running from the threat area to the edge during exploration: blocking edge-vector runs would prevent an SR or FR from predicting that the threat area leads to future occupancy at the obstacle edge, thereby disrupting their ability to identify a subgoal there. A predictive map would also account for the result that mice are slow to update their escape routes after an obstacle is removed but update rapidly when the shelter is moved (Shamash et al. 2021): with the SR, changes to the environment are learned gradually, whereas changes to the reward structure are incorporated immediately (Russek et al. 2017). Our results thus add to recent work showing that predictive maps explain spatial cognition better than purely model-free or model-based mechanisms (de Cothi et al. 2021; Stachenfeld et al. 2017).

Standard models of predictive mapping in RL would nonetheless require several extensions in order to capture our results. One clear requirement would be to abstract beyond elemental actions (e.g., the usual *go 1 unit up, down, left, or right*) by adding the capacity to learn subgoals, as in the option-learning framework (Sutton et al. 1999). Enticingly, neural signatures of option learning have been found in humans (Ribas-Fernandes et al. 2011), but this line of work has not yet been extended to rodent models. Second, our previous results showed that subgoal learning is dramatically reduced when the lights are turned off, when the obstacle is a hole instead of a wall, and when the shelter is not present during the exploration period (Shamash et al. 2021). Thus, the landmark-guided and goal-directed nature of edge-vector runs appears to be important for rapidly learning subgoals in a totally novel environment. This aligns with results in RL showing that options can be learned from successful, intrinsically motivated actions (Barto et al. 2004) and that the rich visual and interactive experiences inherent to biological learning could be essential for mimicking animals’ cognitive abilities *in silico* (Hill et al. 2020). Finally, if mice were limited to a predictive map, we would expect the initiation set (locations from which subgoal escapes are initiated) to be biased toward the locations that predict future occupancy at the obstacle edge, such as edge-vector run start points or anywhere along the obstacle. Instead, we found that mice possess a spatial awareness of distances within the environment that goes beyond their exact history of movements. This suggests a collaboration between high-level policy generation with a predictive representation and fine-grained planning with information from a Euclidean cognitive map. This combination could be instantiated by building a predictive map with input from cells that encode Euclidean distance information, such as object-vector cells (de Cothi and Barry 2020) or grid cells (Banino et al. 2018), or by having both a predictive place-cell map and a Euclidean grid-cell map that share control over behavior (similar to Edvardsen et al. 2020).

Though lacking the formal precision of reinforcement learning, sensorimotor enactivism provides a complementary perspective for understanding how action-driven learning could catalyze a map-based planning process. Sensorimotor enactivism is a strain of research in the cognitive sciences that emphasizes the importance of specific motor actions and their sensory consequences in facilitating a wide array of cognitive feats (Clark 1999; Mataric 1992; Ward et al. 2017). For example, Ballard et al. 1997 suggest that saccadic eye movements act as visual ‘pointers’ which bind external objects to ongoing cognitive processes related to those objects, such as planning chess moves (Chase and Simon 1973). These saccades ease the burdens of attentional selection and working-memory demands. In our case, the ‘sensorimotor primitives’ in question are intrinsically motivated runs to a visually salient edge, and their function would be to refine the computational search for important locations or compartments within the environment. Thus, rather than mentally searching for optimal subgoals in an internal spatial representation, mice enact a search algorithm. Running from the threat area to the obstacle edge triggers the mouse to notice that this location provides special access to the shelter. This new inference could interface with a classic hippocampal cognitive map, embedding a subgoal location within it. Thenceforth, the mouse would use its subgoal memory whenever located beyond the limits of where a sensorimotor strategy (e.g. visual or tactile guidance past the obstacle) would operate effectively; this could underlie the spatial specificity of the initiation set. Finally, once the obstacle is removed, this subgoal policy could remain in place until another sensorimotor insight alerts the mouse to the presence of a shortcut.

A key remaining question is to define the scope of this action-driven mapping process and its relationship to classical map-based cognition: does experience with action-driven mapping lay the foundation for action-independent cognitive mapping, such as the ability to compute subgoals without relying on practice runs? Does action-driven mapping imply a tight coordination between the hippocampal-map and striatal-action circuits often described as competing for control of behavior? Or are these simply independent strategies, deployed in distinct timescales, spatial scales, tasks, brain regions and species? Future work across different species and behaviors will be needed in order to build a broader picture of the role of action-driven mapping in mammalian cognition at large.

## Acknowledgements

This work was funded by a Wellcome Senior Research Fellowship (214352/Z/18/Z) and by the Sainsbury Wellcome Centre Core Grant from the Gatsby Charitable Foundation and Wellcome (090843/F/09/Z) (T.B.) and the Sainsbury Wellcome Centre PhD Programme (P.S.). We thank members of the Branco lab for discussions; Jesse Geerts, Marcus Stephenson-Jones, Caswell Barry, Catarina Albergaria and Ted Moscovitz for comments on the manuscript; Panagiota Iordanidou for experimental support; the Sainsbury Wellcome Centre Neurobiological Research Facility and FabLabs for technical support.

## Methods

**Table 1:** List of all experiments

| ID | Experimental setup | M2 stimulation | Mice | Figures |
| --- | --- | --- | --- | --- |
| 1 | Obstacle removal | Injection/implantation, no stim | 8 | F1,2, SF3,4,7 |
| 2 | Obstacle removal | Stop edge-vector runs | 8 | F1,2, SF1,3,4 |
| 3 | Open field–no obstacle | Injection/implantation, no stim | 8 | F2, SF4 |
| 4 | Obstacle removal | Stop edge-vector runs after two | 8 | SF5,6,7 |
| 5 | Obstacle removal | Stop edge-to-shelter runs | 8 | F3, SF6,7 |
| 6 | Obstacle removal | Stop threat-area-to-left-side runs | 8 | F4, SF6,7 |
| 7 | Obstacle removal–threat zone II | None | 8 | SF7 |
| 8 | Obstacle removal–threat zone III | None | 8 | F4, SF7 |
| 9 | Two-chamber place preference | Paired with one chamber | 8 | SF1 |
| 10 | Open field–no obstacle or shelter | Test effects of three laser powers | 4 | SF1 |

### Animals

All experiments were performed under the UK Animals (Scientific Procedures) Act of 1986 (PPL 70/7652) after local ethical approval by the Sainsbury Wellcome Centre Animal Welfare Ethical Review Body. We used 36 singly housed (starting from 8 weeks old), male, 8–12-week-old C57BL/6J mice (Charles River Laboratories) during the light phase of the 12-h light/dark cycle. Mice were housed at 22°C and in 55% relative humidity with ad libitum access to food and water.

#### Re-use over multiple sessions

For the exploration + escape experiments in implanted mice (experiments 1-6): four of the eight mice were naive, and this was their first behavioral session of any sort. The remaining four mice had experienced a previous session 5-7 days prior. Their previous session was not allowed to be the same exact experiment as the second session but was otherwise selected randomly. The effects of having a previous session on escape behavior were modest (Supp. Fig. 4c-d), and do not impact the interpretation of our results. For the place-preference experiment and laser-power test, mice were randomly selected from those that had already experienced their behavioral sessions in experiments 1-6. For the experiments in unimplanted mice, experiment #7 was performed in naive mice, and experiment #8 was performed 5-7 days later, with the same set of mice.

#### Exclusion criteria

Data from mice with zero escapes in the session (three mice: due to staying in the shelter; two mice: due to not responding to the threat stimulus; one mouse: due to climbing down from the platform; all mice had a previous session) were excluded, and a replacement session was performed 5-7 days later in a randomly selected mouse.

### Viral injection and fiber-optic cannula implantation

#### Surgical procedure

Mice were anaesthetized with isoflurane (5%) and secured on a stereotaxic frame (Kopf Instruments). Meloxicam was administered subcutaneously for analgesia. Isoflurane (1.5–2.5% in oxygen, 1 l min^−1^) was used to maintain anesthesia. Craniotomies were made using a 0.7 mm burr (Meisinger) on a micromotor drill (L12M, Osada), and coordinates were measured from bregma. Viral vectors were delivered using pulled glass pipettes (10 *µ*l Wiretrol II pulled with a Sutter-97) and an injection system coupled to a hydraulic micromanipulator (Narishige), at approximately 100 nl min^−^1. Implants were affixed using light-cured dental cement (3M) and the surgical wound was closed using surgical glue (Vetbond).

#### Injection and implantation

Mice were injected with 120 nL of AAV9/CamKIIa-ChR2-EGFP in the right, anterior premotor cortex (AP: 2.4 mm, ML: 1.0 mm, DV: -0.75 mm relative to brain surface) and implanted with a magnetic fiber-optic cannula directly above the viral injection (DV: -0.5 mm) (MFC_200/245-0.37_1.5mm_SMR_FLT, Doric). All behavioral sessions took place 2-4 weeks after the injection/implantation.

#### Histology

To confirm injection and implantation sites, mice were terminally anaesthetized by pentobarbital injection and decapitated for brain extraction. The brains were left in 4% PFA overnight at 4°C. 100um-thick coronal slices were acquired using a standard vibratome (Leica). The sections were then counter-stained with 4’,6-diamidino-2-phenylindole (DAPI; 3 *µ*M in PBS), and mounted on slides in SlowFade Gold antifade mountant (Thermo Fisher, S36936) before imaging (Zeiss Axio Imager 2). Histological slice images were registered to the Allen Mouse Brain Atlas (Allen Institute for Brain Science 2015) using SHARP-Track (Shamash et al. 2018), to find the fiber tip coordinates.

### Behavioral apparatus

#### Platform and shelter

Experiments took place on an elevated white 5-mm-thick acrylic circular platform 92 cm in diameter. The platform had a 50×10 cm rectangular gap in its center. For conditions with no obstacle (all post-exploration escapes and the entirety of experiments 3 and 10), this was filled with a 50×10 cm white 5-mm-thick acrylic rectangular panel (Supp. Fig. 1b). For conditions with the obstacle present (the exploration period in experiments 1-2 and 4-8), this was filled with an identical panel that, attached to an obstacle: a 50 cm long x 12.5 cm tall x 5 mm thick white acrylic panel (Supp. Fig. 1a). The shelter (Supp. Fig. 1) was 20 cm wide x 10 cm deep x 15 cm tall and made of 5-mm-thick transparent red acrylic, which is opaque to the mouse but transparent to an infrared-detecting camera. The shelter had a 9cm-wide entrance at the front, which extended up to the top of the shelter and then 5 cm along its ceiling; this extension of the opening allowed the optic fiber, which was plugged into the mouse’s head, to enter the shelter without twisting or giving resistive force.

#### Additional hardware

The elevated platform was located in a 160 cm wide x 190 cm tall x 165 cm deep sound-proof box. A square-shaped projector screen (Xerox) was located above the platform. This screen was illuminated in uniform, gray light at 5.2 cd m^−^2 using a projector (BenQ). Behavioral session were recorded with an overhead GigE camera (Basler) with a near-infrared selective filter, at 40 frames per second. Six infrared LED illuminators (TV6700, Abus) distributed above the platform illuminated it for infrared video recording. All signals and stimuli, including each camera frame, were triggered and synchronized using hardware-time signals controlled with a PCIe-6351 and USB-6343 input/output board (National Instruments), operating at 10 kHz. The platform and shelter were cleaned with 70% ethanol after each session.

#### Data acquisition software and online video tracking

Data acquisition was performed using custom software in the visual reactive programming language Bonsai (Lopes et al. 2015). In order to automatically deliver laser and auditory stimuli (see below), mice were tracked online during each behavioral session. Online tracking was based on the mouse being darker than the white acrylic platform; we used the following Bonsai functions, in this order: BackgroundSubtraction, FindContours, BinaryRegionAnalysis, and LargestBinaryRegion.

### Closed-loop optogenetic stimulation

Laser stimuli consisted of 2-sec, 20-HZ square-wave pulses at 30 mW (duty cycle 50%, so 15 mW average power over the two seconds) supplied by a 473-nm laser (Stradus 472, Vortran). For experiment #5, we instead used 5-sec pulses. The laser was controlled by an analog signal from our input/output board into the laser control box. At the beginning of each session, the mouse was placed in an open 10×10 cm box and the magnetic fiber-optic cannula was manually attached to a fiber-optic cable (MFP_200/230/900_0.37_1.3m_FC-SMC, Doric). A rotary joint (Doric) was used to prevent the cable from twisting. Finally, the rotary joint was connected to the laser via a 200-*µ*m core patch cable (ThorLabs).

At the beginning of each mouse’s first session, the mouse was placed in a 10×10 cm box, and two 2-sec stimuli were applied. If these did not evoke stopping and leftward turning (2/24 mice), then the mouse was assigned to one of the laser-off conditions (experiment 1 or 3). During laser-on sessions, the criteria for triggering laser stimuli were: 1) the mouse crosses the ‘trip wire’ (illustrated in Figure 1, 3, 4); and 2) the mouse is moving in the ‘correct’ direction. For blocking edge-vector and edge-to-shelter runs, the direction was determined by a directional speed threshold: moving toward the shelter area (i.e., south) at > 5 cm sec^−^1. For blocking threat-zone-to-left-side runs, mice had to be moving toward the left side (i.e., west) at > 5 cm sec^−^1. These speed thresholds are low enough to be effective at catching all cases in which the mouse crosses the trip wire in a particular direction. These criteria were computed online using the Bonsai software described in the previous section. The laser pulses were emitted with a delay of 300-400 ms after being triggered. Up to three subsequent 2-sec pulses (or one 5-sec pulse in experiment #5) were triggered manually if the mouse continued moving forward.

Mice usually took 1-3 minutes to enter the shelter for the first time, and these first minute(s) of exploration typically contains relatively vigorous running. Since subgoal learning does not occur in this setting without a shelter in the environment (Shamash et al. 2021), the laser-on condition was initiated only after the mouse entered the shelter for the first time.

### Exploration and Escape behavior

#### Auditory threat stimuli

Threat stimuli were loud (84 dB), unexpected crashing sounds played from a speaker located 1 m above the center of the platform (Supplementary Audio 1 and 2). Sounds (‘smashing’ and ‘crackling fireplace’) were downloaded from soundbible.com. They were then edited using Audacity 2.3.0, such that they were 1.5 sec long and continuously loud. Stimuli alternated between the ‘smashing’ sound and the ‘crackling’ sound each trial, to prevent stimulus habituation. The volume was increased by 2 dB after time a stimulus failed to elicit an escape, up to a maximum of 88 dB. When a threat trial began, the stimuli repeated until the mouse reached the shelter or for a maximum of 9 secs.

#### Triggering escapes

The criteria for activating a threat stimulus were 1) the mouse is currently in the threat zone (illustrated in Figure 2);2) the mouse was in the threat zone 1.5 seconds ago; 3) the mouse is moving away from the shelter at >5 cm s^−^1 (this ensures that escape runs are always initiated after the stimulus onset); 4) the most recent threat stimulus occurred >45 sec ago. These criteria were computed online using the Bonsai software described above, and auditory threat stimuli played automatically when all four criteria were met. Experiments were terminated after six successful escapes or one hour. In experiments 7-9, criterion #2 was not applied. For experiment #8, experiments were terminated after ten escapes rather than six, as this threat zone allowed for more trials. Reaching the shelter was defined as reaching any point within 10 cm of the shelter entrance, and escapes were considered successful if they reached the shelter within the 9-sec stimulus period.

#### Obstacle removal

After 20 minutes of exploration were complete, as soon as the mouse entered the shelter, the experimenter quickly and quietly removed the central panel containing the obstacle and replaced it with the flat 50×10 cm panel. Mice were then allowed to freely explore and (and trigger escapes) in this open-field platform.

#### Adding bedding to the platform

Bedding from the mouse’s home cage was added to the platform in order to encourage exploration, rather than staying in the shelter throughout the experiment. One pinch (1 gram) of bedding was added to the center of the threat zone in all experiments when either of the following two criteria was met: 1) The mouse did not leave the shelter for five minutes; or 2) The mouse did not enter the threat zone for ten minutes. In order to encourage occupancy of the areas from which edge-vector runs initiate, a pinch of bedding was placed on the left side of the threat zone in experiments #4 and 6, and the left and right sides in experiments 7-8. In order to maintain comparability across conditions, a pinch of bedding was also placed in the same location for the mice with a previous session in experiment #2.

### Place preference assay

Mice were hooked up to the optic fiber as described above and placed into a two-chamber place-preference arena. The arena was made of 5-mm-thick transparent red acrylic (opaque to the mouse) and consisted of two 18 cm long x 18 cm wide x 18 cm tall chambers connected by a 8cm-long opening. To make the chambers visually distinguishable, one chamber had a 10×10 cm x-shaped white acrylic piece affixed to its back wall and the other had a filled-in, 10cm-diameter circular white acrylic piece affixed to its back wall. The stimulation chamber (left or right) was pseudoramdomly determined before each session, such that both sides ended up with four mice. After a 1-min habituation period, a series of four 2-sec laser stimuli were manually triggered whenever the mouse fully entered the stimulation chamber. A minimum of one minute was given in between each trial, and a total of six stimulation series were delivered. After the last stimulation, one minute was given so that the occupancy data would not be biased by always starting in the stimulation chamber. Then, the next 20 minutes were examined to test for place aversion in the stimulation chamber. This assay is adapted from the conditioned place preference assay (Stamatakis and Stuber 2012) and the passive place avoidance assay (Schlesinger et al. 1983), such that it matches the conditions of our exploration/escape assay (i.e., to be relevant, place aversion must be elicited during the same session as the laser stimulation, and it must be expressed through biases in occupancy patterns)

### Analysis

All analysis was done using custom software written in Python 3.8 as well as open-source libraries, notably NumPy, OpenCV, Matplotlib and DeepLabCut.

#### Video tracking

Video recording was performed with custom software in Bonsai. We used DeepLab-Cut (Mathis et al. 2018) to track the mouse from the video, after labeling 412 frames with 13 body parts: snout, left eye, right eye, left ear, neck, right ear, left upper limb, upper back, right upper limb, left hind limb, lower back, right hind limb and tail base (Video 5). Post-processing includes removing low-confidence tracking, using a median filter with a width of 7 frames and applying an affine transformation to the tracked coordinates to match the common coordinate framework. Videos were generated using custom Python code, the OpenCV library and Adobe AfterEffects.

#### Calculating position, speed and heading direction

For analysis of escape trajectories and exploration, we used the average of all 13 tracked points, which we found to be more stable and consistent than any individual point. To calculate speed, we smoothed the raw frame-by-frame speed with a Gaussian filter (*σ* = 4 frames = 100 ms). To calculate the mouse’s body direction, we computed the vector between the lower body (averaging the lower left limb, lower right limb, lower back, and tail base) and the front of the body (averaging the upper left limb, upper right limb, and upper back). See Video 5 for a visualization of the tracking and of these calculations.

#### Analysis of escape trajectories

The escape target score was computed by taking the vector from the mouse’s position at escape initiation to its position when it was 10 cm in front of the obstacle. Vectors aimed directly at the shelter received a value of 0; those aimed at the obstacle edge received a value of 1.0; a vector halfway between these would score 0.5; and a vector that points beyond the edge would receive a value greater than 1.0. The formula is:

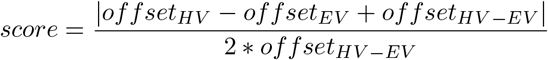

Offset_*HV*_ is the distance from the mouse to where the mouse would be if it took the homing vector; offset_*EV*_ is the distance from the mouse to where the mouse would be if it took the obstacle edge vector; and offset_*HV*_− _*EV*_ is the distance from the homing vector path to the obstacle edge vector path. The threshold for classifying a trajectory as an edge vector (scores above 0.65) was taken from Shamash et al. 2021, where it represented the 95th percentile of escapes in the open-field condition. Escapes with scores under 0.65 were designated as homing vectors. When escape trajectories are limited to escapes on the left side, this refers to escapes that are on the left half of the arena when they cross the center of the platform along the vertical (threat-shelter) axis.

#### The escape initiation point

occurs when mice surpass a speed of 20 cm s^−1^, relative to (i.e., getting closer to) the shelter location. This threshold is high enough to correctly reject non-escape locomotion bouts along the perimeter of the platform but also low enough to identify the beginning of the escape trajectory.

#### Extraction of spontaneous homing runs and edge-vector runs

Homing runs are continuous turn-and-run movements from the threat area toward the shelter and/or obstacle edges. As in Shamash et al. 2021, they are extracted by (1) computing the mouse’s ‘homing speed’ (that is, speed with respect to the shelter or obstacle edges with Gaussian smoothing (/sigma = 0.5 s)) and the mouse’s ‘angular homing speed’ (the rate of change of heading direction with respect to the shelter or obstacle edges); identifying all frames in which the mouse has a homing speed of >15 cm s^−1^ or is turning toward the shelter at an angular speed of >90° per sec; (3) selecting all frames within 1 s of these frames, to include individual frames that might be part of the same homing movement but do not meet the speed criteria; (4) rejecting all frames in which the mouse is not approaching or turning toward an edge or the shelter; and (5) rejecting sequences that take less than one sec or do not decrease the distance to the shelter by at least 20%. Each series of frames that meet these criteria represents one homing run. We limited analysis to he homing runs that started within the threat area (Figure 1a). *Edge-vector runs* are homing runs that enter anywhere within the 10-cm-long (along the axis parallel to the obstacle) x 5-cm-wide (along the axis perpendicular to the obstacle) rectangle centered 2.5 cm to the left of the obstacle edge.

#### Initiation set analysis: logistic regression

Our logistic regression analysis tests the strength of the linear relationship between each spatial metric and the log odds of performing an edge-vector escape. No regularization penalty was used. The strength of the fit was measured using McFadden’s pseudo-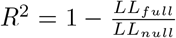, where *LL*_*full*_ is the log likelihood of the logistic regression model fitted with the predictor data and *LL*_*null*_ is the log likelihood of the logistic regression fitted with only an intercept and no predictor data. Pseudo-R^2^ values of 0.2-0.4 represent “excellent fit”(McFadden 1977). To test statistical significance of these values, we performed a permutation test, based on the distribution of pseudo-R^2^ for the same predictor value, across 10,000 random shuffles of the escape responses (edge vector or homing vector).

#### Initiation set analysis: normalizing a metric

To normalize a spatial metric (y, e.g. distance from the center of the arena along the left-right axis) by another metric (x, e.g. distance from the shelter), we computed a linear regression on these variables. We then took the residuals of this prediction (*residual* = *y −ŷ*, where *ŷ* = *slope× x* + *offset*) and correlated them with proportion of edge-vector escapes in each bin. This tells us whether, at a given distance from the shelter, there is still a correlation with distance from the center.

#### Initiation set analysis: correlation analysis

To better visualize the relationship between the mouse’s initial position and the likelihood of executing an edge-vector escape, we binned the spatial metric and computed the correlation to the proportion of edge-vectors in each bin. The widest possible range of values was selected, given the constraints that this range starts and ends on a multiple of 2.5 cm and that all bins contain at least six escapes. From this range, seven equal-sized bins were used. The correlation results were robust to the number of bins used.

#### Statistics

For comparisons between groups, we used a permutation test with the test statistic being the pooled group mean difference. The condition of each mouse (e.g., laser-on vs. laser-off) is randomly shuffled 10,000 times to generate a null distribution and a p-value. We used this test because it combines two advantages: 1) Having the test statistic as the pooled group mean gives weight to each trial rather than collapsing each animal’s data into its mean (as in the t-test or the Mann–Whitney test); 2) It is non-parametric and does not assume Gaussian noise (unlike the repeated-measures ANOVA), in line with much of our data. Tests for increases or decreases (e.g., whether exploration decreased due to laser stimulation) were one tailed. The Wilcoxon signed-rank test was used for the place-preference assay to test whether occupancy in the stimulation chamber was less than 50%. The sample size of our experiments (n=8 mice) was selected based on a power analysis based on the data from Shamash et al. 2021 and a minimum power of 0.8. Ranges in box plots are limited from the first quartile minus 1.5 x IQR to the third quartile plus 1.5 x IQR. Statistically significant results are indicated in the figures using the convention *n.s*.: *p>0.05, *: p<0.05, **: p<0.01 and ***: p<0.001*.

### Data and software availability

The data-acquisition software is available from https://github.com/philshams/bonsai-behavior, and the data-analysis software is available from https://github.com/philshams/behavior-opto-analysis. The data from this study will be made available upon publication.

## Supplementary Figures

**Supplementary Figure 1:**
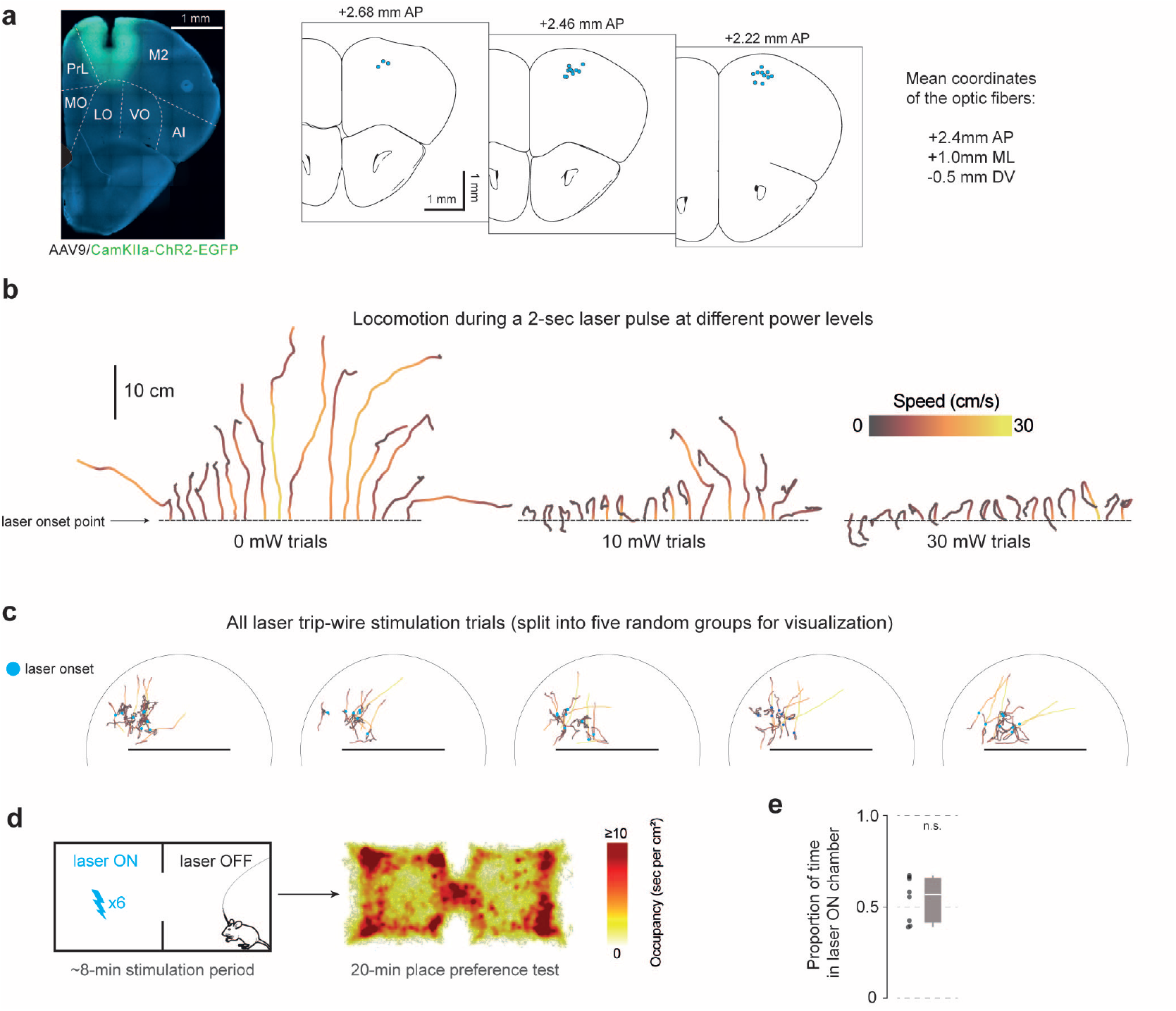
Optogenetic stimulation of right premotor cortex **(a)** Left: example viral injection and optic fiber implantation site. M2: supplementary motor cortex (premotor cortex), PrL: prelimbic cortex, MO/LO/VO: medial/lateral/ventral orbital cortex, AI: agranular insular cortex. Right: Putative optic fiber tip locations are overlaid on brain-slice diagrams adapted from Paxinos and Franklin 2019. Histological slice images were registered to the Allen Mouse Brain Atlas (Allen Institute for Brain Science 2015) using SHARP-Track (Shamash et al. 2018) to find the fiber tip coordinates. The site of injection with channelrhodopsin was 0.25 mm below (ventral to) the fiber tip. AP and ML coordinates are relative to bregma, and DV coordinates are relative to the brain surface. **(b)** Locomotion following a 2-sec, 20-Hz, 30-mW pulse wave (duty cycle 50%) of 473-nm light in implanted mice. Laser stimulation was triggered manually upon initiation of a running bout, in the behavioral platform with no obstacle and no shelter. Each mouse received 4 trials at each laser power, sequentially interleaved. n = 4 mice. Lines are ordered by the distance and direction of movement following laser onset. **(c)** Trajectories before and after laser stimulation, for the edge-vector blocking protocol. n = 8 mice, 3.5 [2.75,6] (median [IQR]) laser stimulation trials per mouse. **(d)** Place preference assay. Each chamber in the place-preference arena (18 cm x 18 cm x 18 cm) has a distinguishing landmark on the back wall (a large cross and a large circle). After a 1-min habituation period, stimulation consisted of six trials of four repeated 2-sec, 20-Hz, 30-mW pulses (24 total pulses). Stimulation was manually triggered when the mouse fully entered the stimulation side, with at least one minute between trials. The side of stimulation was pseudo-randomly selected such that half of the mice were stimulated on each side. For the occupancy heatmap, stimulation is shown as if it were on the left side for all mice. The heatmaps was smoothed with a gaussian filter (*σ* = 0.3 cm). **(e)** Occupancy in the stimulation chamber is not significantly below 50%. p = 0.7, one-tailed Wilcoxon signed-rank test. n = 8 mice.

**Supplementary Figure 2:**
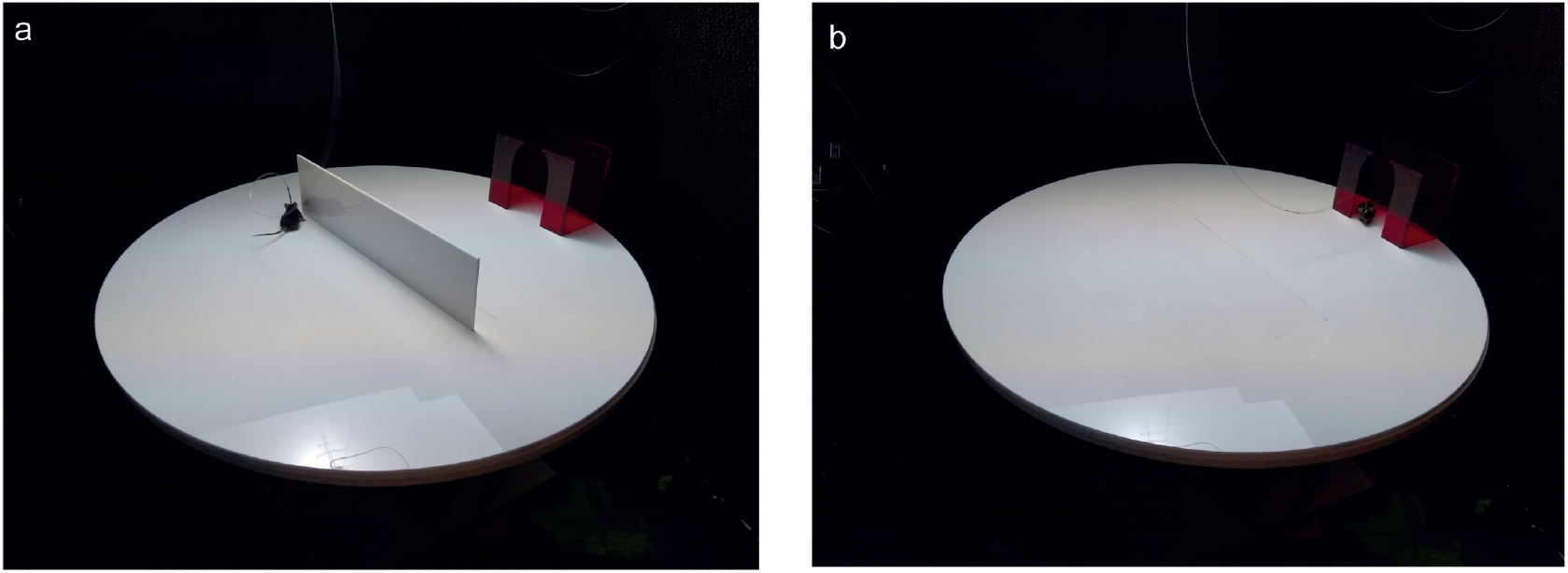
Behavioral platform with and without the obstacle **(a)** The platform with the wall obstacle. The platform is 92 cm in diameter, and the wall obstacle is 50 cm long x 12.5 cm tall. The shelter is 20 cm wide x 10 cm deep x 15 cm tall. It is made from red acrylic that is opaque to the mouse but transparent to red and infrared light. The mouse has just run to the right obstacle edge. **(b)** The platform with no obstacle. A central panel (50 cm wide x 10 cm wide) with the obstacle has been replaced, and a flat panel has been slotted in, in its place. The mouse is sitting in the shelter.

**Supplementary Figure 3:**
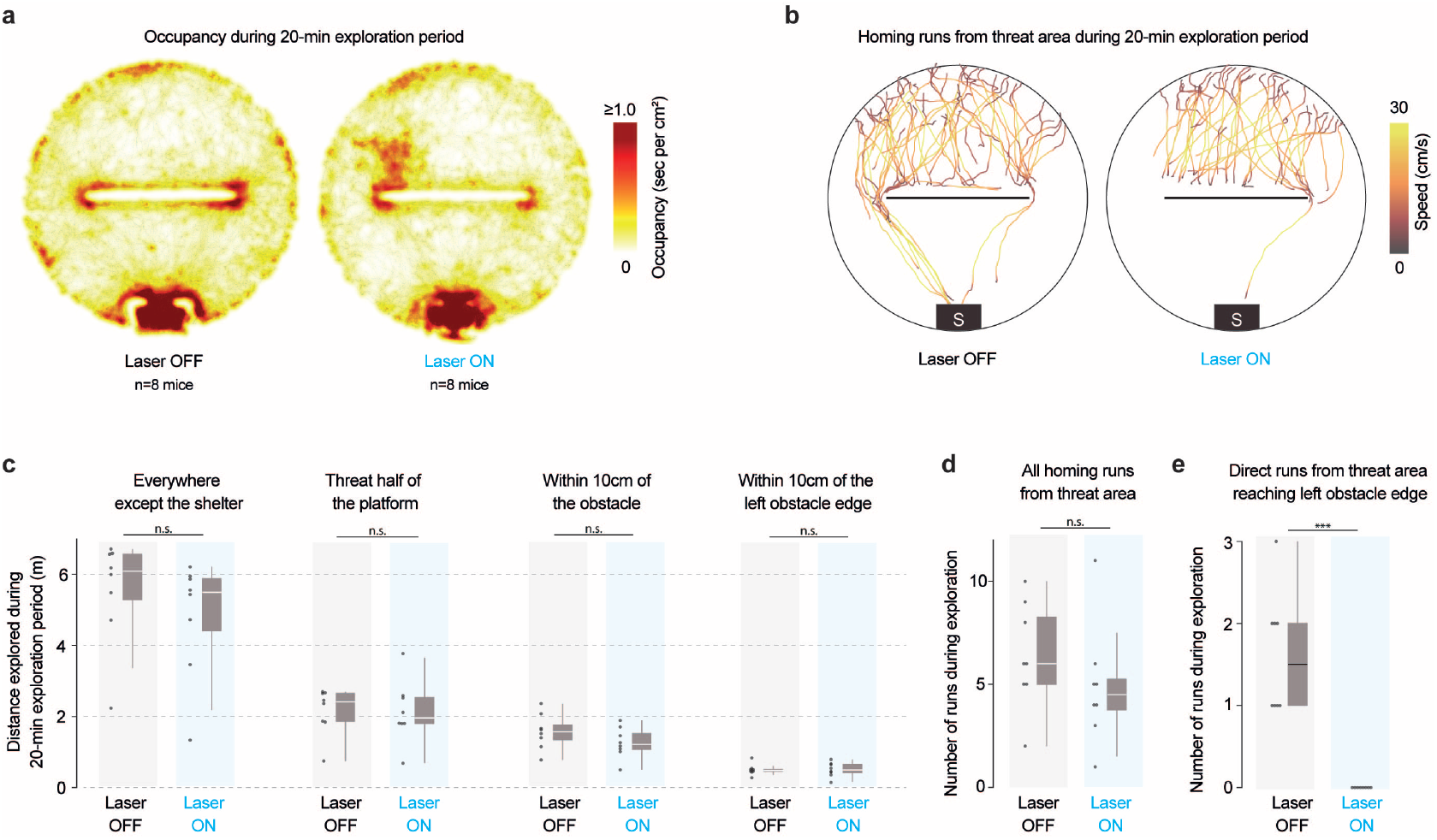
Effect of optogenetic stimulation on exploration **(a)** Occupancy heatmaps are smoothed with a gaussian filter (*σ* = 1 cm). **(b)** Runs from all eight mice in each condition. Left: same runs as in Figure 1a, except with non edge-vector runs also included here. Right: Homing runs do not reach the left obstacle edge due to the closed-loop optogenetic stimulation. **(c)** Distance explored is used instead of time explored to account specifically for active exploration, but the results look similar when time explored is used. Everywhere except the shelter: p = 0.2; threat half: p = 0.5; obstacle: p = 0.1, edge: p = 0.5, one-tailed permutation tests. **(d)** Total number of homing runs (trajectories shown in panel b): p = 0.15. **(e)** Runs reaching the left edge: p = 3 *×* 10^−5^, one-tailed permutation tests.

**Supplementary Figure 4:**
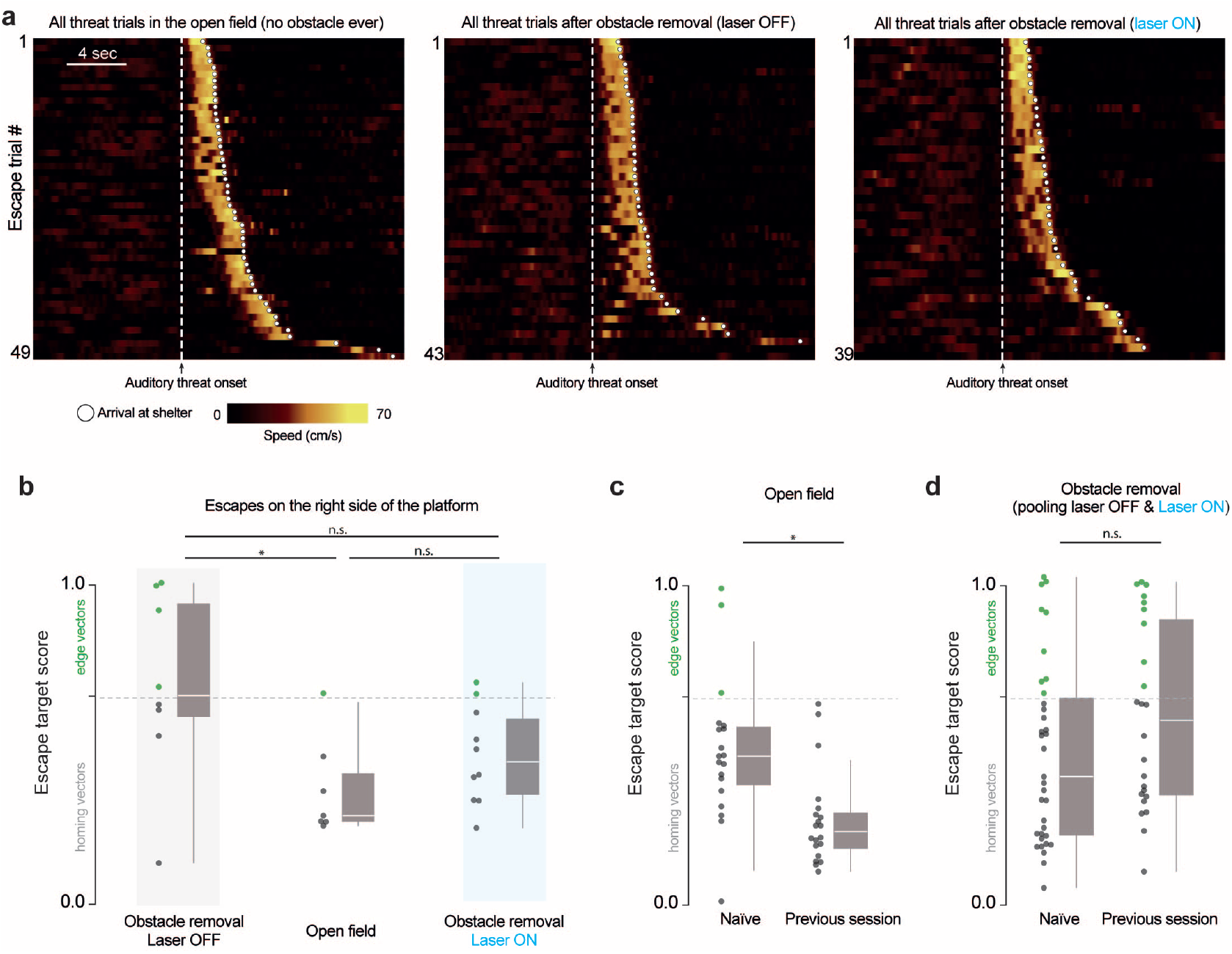
Escape behavior **(a)** Open field: n = 8 mice, 29 trials; obstacle removal (laser off): n = 8 mice, 26 trials; obstacle removal (laser on): n = 8 mice, 23 trials. Speed is smoothed with a gaussian filter (*σ* = 100 ms). **(b)** Escapes on the right side are defined as escapes that, upon passing the center of the arena along the shelter-threat (north-south) axis, were on the right half of the platform. In the main figures, we limited analysis to escapes on the left side of the platform (measured at the center of the shelter-threat axis). This allows us to evaluate our model, in which edge-vector runs generate subgoals one at a time, at the targeted edge. We observed that there were fewer escapes on the right side across all conditions, possibly due to an environmental bias or an effect of the brain implant. As a result, there is not enough data determine whether the laser manipulation has an effect on right-side escapes. Obstacle removal (laser off) vs. open field: p = .045; Obstacle removal (laser on) vs. open field: p = .1; Obstacle removal (laser off) vs. obstacle removal (laser on): p = .1, one-tailed permutation tests on proportion of edge-vector escapes. Open field: 10 escapes; Obstacle removal (laser off): 8 escapes; Obstacle removal (laser on): 10 escapes. **(c)** In each experiment, 4/8 mice were naive and 4/8 mice had had a previous behavioral session, in a random condition. Mice with a previous session targeted the shelter more accurately in the open-field environment. p = .03, one-tailed permutation test on proportion of edge-vector escapes. **(d)** Mice with a previous session did not execute significantly more edge-vector escapes than naive mice. p = 0.2, one-tailed permutation test on proportion of edge-vector escapes.

**Supplementary Figure 5:**
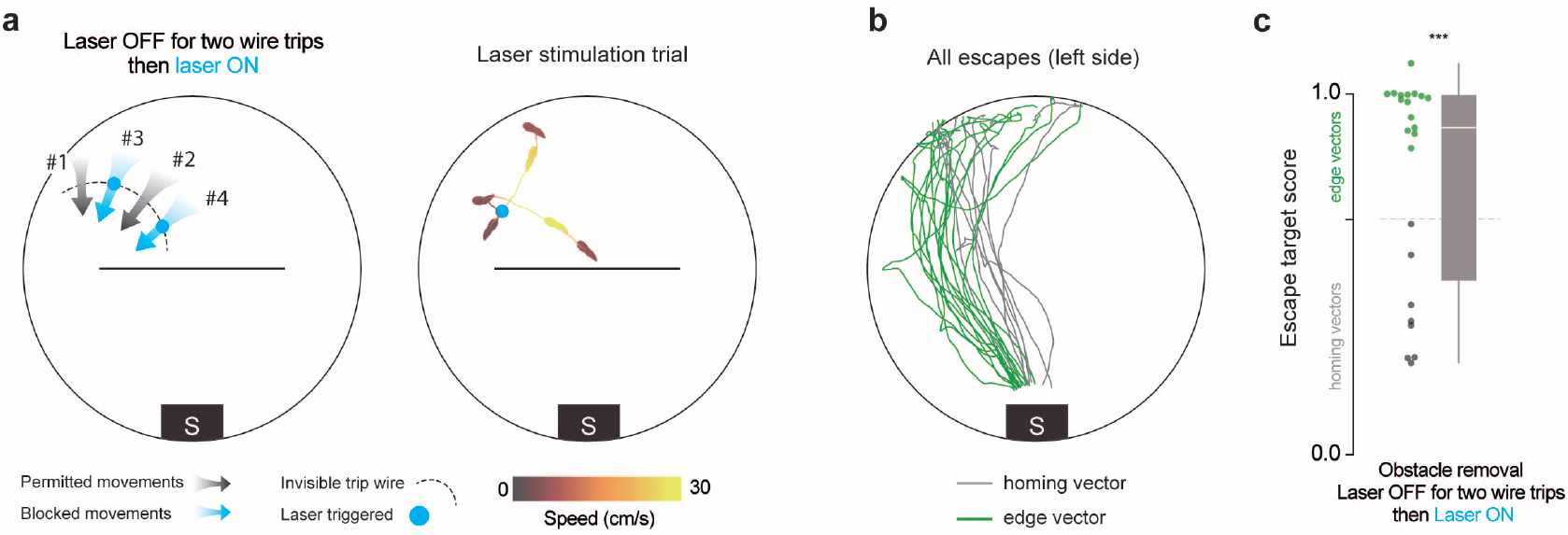
Blocking edge-vector runs after allowing two does not reduce subgoal learning **(a)** Schematic of stimulation blocking all but the first two edge-vector runs. Stimulation protocol follows the edge-vector-blocking protocol from figure 1, except that the first two trip-wire crossings are allowed to occur with no laser stimulation. This entailed a median of 3.0 laser-stimulation trials per mouse, compared to 3.5 in the original experiment from figures 1-2. To make sure these numbers would be comparable, we placed a pinch of bedding on the left side of the threat area, which encouraged the mice to travel to and from that area (see Methods). The example shows four seconds after laser onset: the mouse was stimulated for two seconds, and then ran toward the center of the obstacle. **(b)** Escapes after obstacle removal. n = 8 mice, 23 escapes. **(c)** Obstacle removal (block after two crossings) vs. open field: p = 3*×* 10^−4^ (***); vs. obstacle removal (block edge vectors): p=.003; vs. obstacle removal (laser off): p = .9, one-tailed permutation tests on proportion of edge-vector escapes.

**Supplementary Figure 6:**
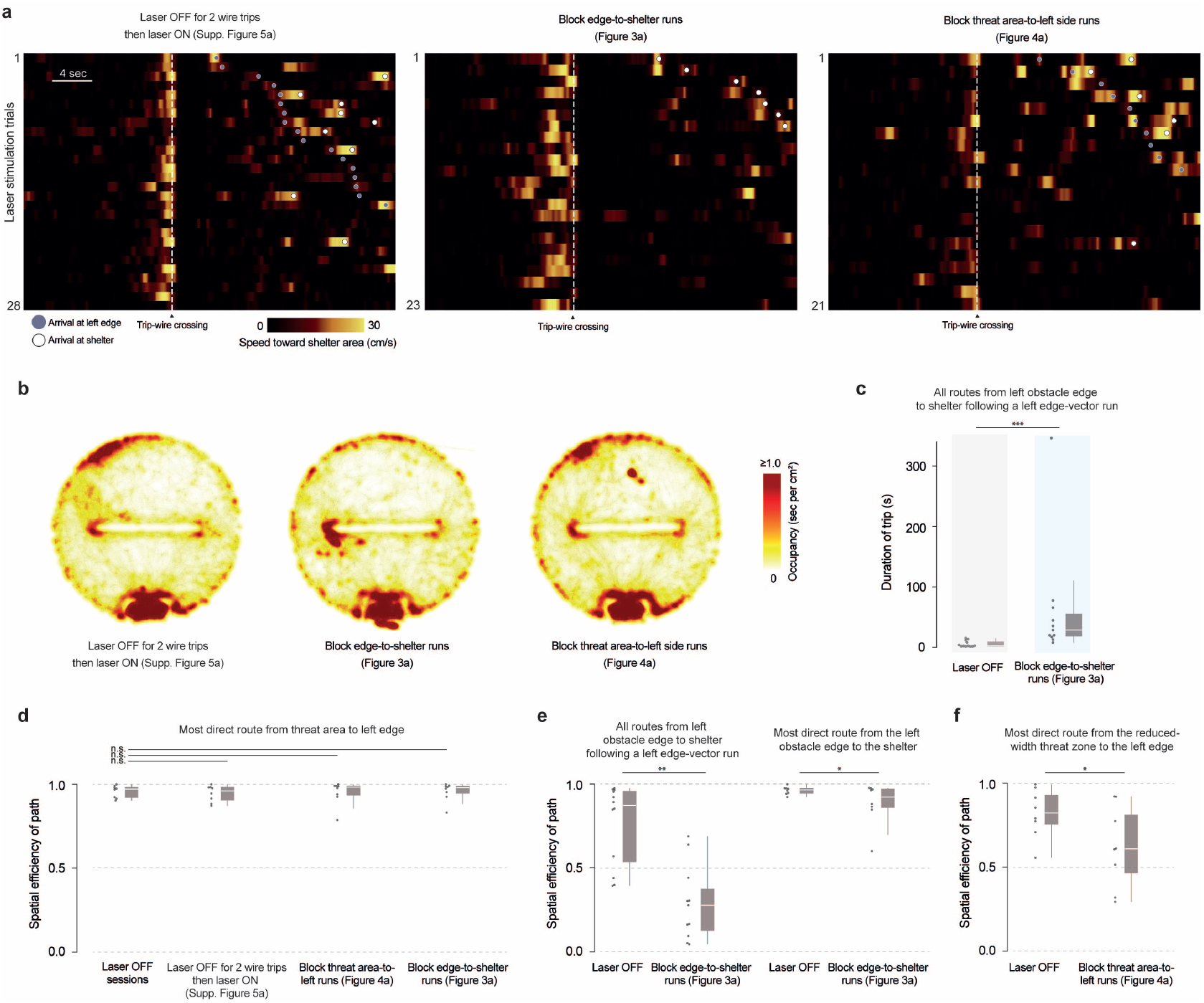
Effects of optogenetic stimulation in the three laser-on control experiments **(a)** All trip-wire crossings, with laser stimulation. Left, center: mice must be moving toward the shelter area (i.e., southward) in order to trigger the trip wire. Right: mice must be moving toward the left side in order to trigger the trip wire. **(b)** Occupancy heatmaps are smoothed with a gaussian filter (*σ* = 1 cm) and overlaid on all movements for all mice (transparent gray dots). There is increased occupancy in the north-west area of the platform in the laser-off-then-on; block threat-to-left-side; and original block-edge-vector conditions due to a pinch of bedding being placed in that area (see Methods). There is increased occupancy near the left obstacle edge in the block edge-to-shelter condition due to the optogenetic stimulation taking place there. **(c)** Each dot represents one trip between the left obstacle edge and the shelter. Laser off median: 2.5 sec; Block edge-to-shelter median: 29 sec. Block edge-to-shelter vs. laser off: p = 0.001, one-tailed permutation test. **(d)** Each dot represents once session (the most efficient path that took place during that session). Laser off-then-on vs. laser off: p = 0.4; block threat-to-left-side vs. laser off: p = 0.3; block edge-to-shelter vs. laser off: p = 0.2, one-tailed permutation tests. **(e)** Left: Each dot represents one trip between the left obstacle edge and the shelter; p = 0.002, one-tailed permutation test. Right: each dot represents one session; p = 0.02, one-tailed permutation test. The effect on the most direct route in the session is relatively weak; this is because, in this experiment, we only blocked edge-to-shelter movements that followed edge-vector runs (i.e., passed the original trip wire from Figure 1). Thus, edge-to-shelter movements that did not pass through the threat area (e.g., running from the shelter to the edge and back) were spared. **(f)** Each dot represents one session; p = 0.02, one-tailed permutation test.

**Supplementary Figure 7:**
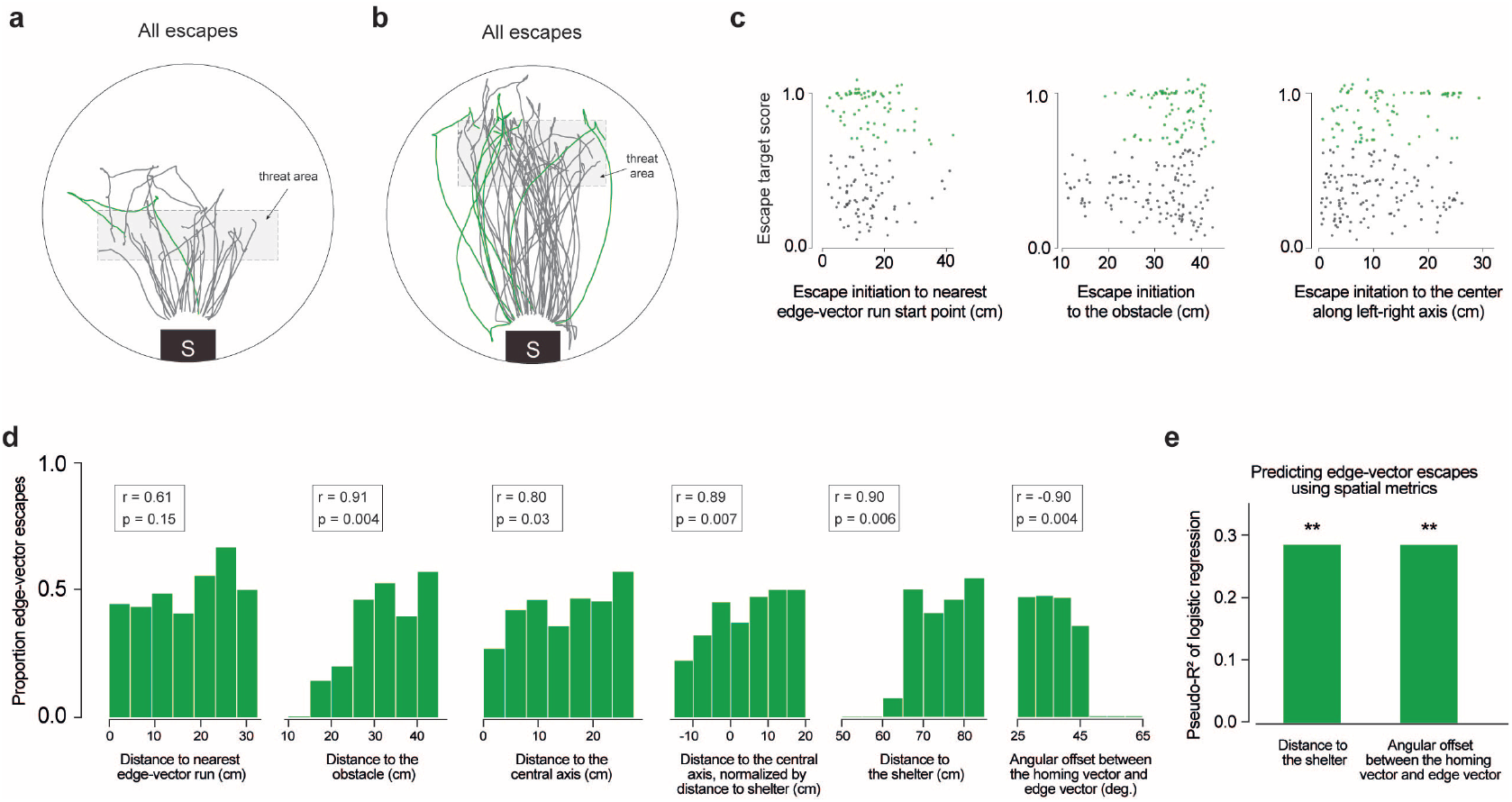
Relationship between escape onset location and subgoal behavior. **(a)** Escapes triggered after obstacle removal, using the new threat zone, indicated by the red dotted lines. Only 1/28 escapes (the green trace) begins by moving toward the obstacle edge location; however, this appears to be a continuation of the pre-threat movement rather than a genuine subgoal escape. **(b)** Escapes triggered after obstacle removal, using another new threat zone, indicated by the red dotted lines. **(c)** Distance metrics plotted against the escape target score. Edge-vector escapes are plotted in green. Distances are measured from the escape initiation point. For the distance to the nearest spontaneous edge-vector run start point (top), only runs toward the same side as the escape are considered. **(d)** To visualize the relationship between position and edge-vector probability, each spatial metric is put into seven equal-sized bins, and the proportion of edge-vector escapes in each bin is taken. All bins have at least six escapes. r-values and p-values come from the Pearson correlation between the spatial bin and the proportion of edge-vector escapes. **(e)** McFadden’s pseudo-R^2^ measures the strength of the relationship between each metric and the odds of executing edge-vector escapes. Distances and angles are measured from the escape initiation point of each escape. For the distance to the nearest spontaneous edge-vector run start point, only runs toward the same side as the escape are considered. Distance to the shelter: pseudo-R^2^=0.29; p=0.006. Angular offset between the homing vector and the edge vector: pseudo-R^2^=0.29; p=0.006. P-values come from a permutation test using 10,000 random shuffles of the edge-vector/homing-vector labels, with the pseudo-R^2^ as the test statistic.

